# Opposing Roles of the Fork-head box genes FoxM1 and FoxA2 in Hepatocellular Carcinoma

**DOI:** 10.1101/383604

**Authors:** Vaibhav Chand, Akshay Pandey, Dragana Kopanja, Grace Guzman, Pradip Raychaudhuri

**Affiliations:** Department of Biochemistry and Molecular Genetics, University of Illinois, College of Medicine, 900 S. Ashland Ave., Chicago, IL-60607.; Department of Pathology, University of Illinois, College of Medicine, Chicago, IL-60612.; Jesse Brown VA Medical Center, 820 S. Damen Ave., Chicago, IL-60612.

**Author notes:** Corresponding Author: Pradip Raychaudhuri. These two authors made equal contribution (Vaibhav Chand, Akshay Pandey).

**Keywords:** DNMT3b, FoxA2, FoxM1, Hepatocellular carcinoma, Retinoblastoma protein

## Abstract

The fork-head box transcription factor FoxM1 is essential for hepatocellular carcinoma (HCC) development and its overexpression coincides with poor prognosis. Here, we show that the mechanisms by which FoxM1 drives HCC progression involve overcoming the inhibitory effects of the liver differentiation gene FoxA2. First, the expression patterns of FoxM1 and FoxA2 in human HCC are opposite. We show that FoxM1 represses expression of FoxA2 in G1 phase, a phase in the cell cycle in which cells can undergo differentiation. Repression of FoxA2 in G1 phase is important, as it is capable of inhibiting expression of the pluripotency genes that are expressed mainly in S/G2 phases. Using a transgenic mouse model for oncogenic Ras-driven HCC, we provide genetic evidence for a repression of FoxA2 by FoxM1. Conversely, FoxA2 inhibits expression of FoxM1, and inhibits FoxM1-induced tumorigenicity of HCC cells. Moreover, expression of FoxA2 in mouse liver expressing activated Ras inhibits FoxM1 expression and inhibits HCC progression. The observations provide strong genetic evidence for an opposing role of FoxM1 and FoxA2 in HCC progression.

**AUTHOR SUMMARY:** Liver cancer remains untreatable because it is diagnosed at a stage when the cancer is aggressive and resistant to therapeutics. The mechanism that drives aggressive liver cancer is poorly understood. These cancers are made up of poorly differentiated cancer cells. Interestingly, the FoxM1 gene is overexpressed in the aggressive liver cancers. Although FoxM1 is important for expression of the proliferation genes, it does not explain why it is overexpressed mainly in the undifferentiated cancers. The current study addresses this puzzle. Our previous studies demonstrated that FoxM1 increases expression of the pluripotency genes that are expressed mainly in the stem-like cells. In the current manuscript we show that, in addition to activating the pluripotency genes, FoxM1 inhibits expression of the liver differentiation gene FoxA2. Overexpression of FoxM1 is important for this inhibition function, as it involves the retinoblastoma family of proteins, which are often inactivated in cancer cells, and thus, are of low-abundance. Moreover, the inhibition of FoxA2 is significant because FoxA2 could inhibit expression of the pluripotency genes as well as FoxM1. The observations provide new insights into how FoxM1 drives progression of aggressive liver cancer.

## INTRODUCTION

Hepatocellular carcinoma (HCC) is the second most fatal malignancies in men worldwide [1, 2]. Development of HCC has been linked to viral hepatitis, alcohol abuse, as well as non-alcoholic steatohepatitis [3-5]. Irrespective of its etiology, one-year survival with intervention is very low. That is partly due to high rate of recurrence of the cancer resulting from intrahepatic and extrahepatic metastasis [6, 7]. Recent studies have linked aggressive progression of HCC to over-expression of the forkhead box transcription factor FoxM1. For example, over-expression of FoxM1 has been shown to strongly correlate with poor prognosis and high-grade progression of HCC [8-10]. Studies with mouse models provided strong causal link between FoxM1 and aggressive progression of HCC. It was shown that FoxM1 is essential for development of HCC in a chemical carcinogenesis model [11]. Deletion of FoxM1 in the adult liver blocked Diethylnitrosamine (DEN)-induced HCC development. Moreover, in the same model of chemical carcinogenesis, deregulated FoxM1 drives highly aggressive, metastatic progression of HCC [12]. In the absence of p19Arf, FoxM1 stimulates all steps of metastatic progression [12]. Consequently, inhibition of FoxM1 impedes metastatic progression of HCC [12].

Similar observations were made with a transgenic mouse model expressing activated HRas in the liver, driven by the albumin promoter [10]. In that model, HRas-induced HCC coincides with an increased expression of FoxM1. Conditional deletion of FoxM1 after HCC development causes inhibition of cancer progression. Hepatic progenitor cells for HCC have been characterized in the chemical carcinogenesis model. They express cell surface markers CD44 and EpCAM [13]. Those cells were detected also in HRas-transgenic mouse model for HCC. They account for about 30 to 40% of the HCC cells in tumor sections [10]. Interestingly, deletion of FoxM1 causes a preferential loss of those cells in the tumor nodules, indicating that FoxM1 is critical for the CD44 +ve and EpCAM +ve HCC cells [10]. CD44 and EpCAM are expressed also by the human hepatic cancer stem cells. Moreover, the hepatic cancer stem cells in human HCC lines are dependent upon FoxM1, as deletion of FoxM1 causes a preferential loss of the cancer stem cells [10]. In that regard, it is noteworthy that FoxM1 is a critical downstream factor of a variety of cancer signaling pathways, including Wnt/b-catenin signaling, that promote cancer stem cells [14].

FoxM1 stimulates expression of the pluripotency genes c-Myc, Oct4, Sox2 and Nanog (reviewed in [15, 16]). In P19 embryonic carcinoma cells, knockdown of FoxM1 causes the cells to undergo differentiation with concomitant loss of expression of the pluripotency genes [15]. Also, expression of FoxM1 induces expansion of human epithelial stem cells and increases expression of the pluripotency genes [17]. In human neuroblastoma cells, depletion of FoxM1 induces expression of differentiation markers with a loss of tumorigenicity and inhibition of the pluripotency genes [18]. Similar results were observed also in HCC. Expression of FoxM1 in human HCC cells, Huh7 and Hep3B, increases expression of the pluripotency genes, whereas knockdown of FoxM1 reduces expression of those genes [10]. Also, in the mouse model of HRas-induced HCC, deletion of FoxM1 causes a reduction in the expression of those genes [10]. It is, therefore, likely that FoxM1 by increasing expression of the pluripotency genes supports high-grade progression of HCC. However, it is unclear whether activation of the pluripotency genes is sufficient to generate undifferentiated cancer cells that drive aggressive progression.

FoxM1 also possesses transcriptional repression activities. In the mammary gland, FoxM1 is expressed at high levels in the stem/progenitor cells to regulate their differentiation [19]. Deletion of FoxM1 decreases the population of the stem/progenitor cells and increases the population of differentiated luminal cells, and the scenario is opposite when FoxM1 is over-expressed in the mammary gland [19]. FoxM1 regulates the luminal differentiation by repressing expression of the luminal transcription factor GATA3. Here we show that the repression function of FoxM1 is critical for aggressive HCC progression.

## RESULTS

### Opposite expression pattern of FoxM1 and FoxA2 in hepatocellular carcinoma

FoxA2 is important for liver development [20]. Interestingly, a recent study indicated that expression of FoxA2 is down-regulated in metastatic HCC [21]. Since deregulation of FoxM1 in HCC drives metastasis [12], we investigated whether FoxM1 and FoxA2 have opposite effects on aggressive progression of HCC. An analysis of the publicly available Roessler’s dataset revealed significant opposite expression patterns of FoxM1- and FoxA2-mRNAs in HCC (supplemental Fig. S1A). However, TCGA dataset did not show significant correlation between RNA expressions of FoxM1 and FoxA2. That could be due to contributions from other cell types known to be present in HCC samples. Therefore, we carried out immunohistochemical staining for the FoxM1 and FoxA2 proteins using tissue microarrays derived from consecutive sections of HCC specimens. There was an obvious difference in the expression pattern of FoxM1 and FoxA2. In the grade I samples, FoxA2 is vividly detectable, whereas the nuclear expression of FoxM1 is low (Fig. 1A-B). On the other hand, in the grade III specimens, expression of FoxA2 is low, but there was abundant expression of FoxM1. Pearson correlation analyses of FoxM1 and FoxA2 expression in the consecutive TMAs indicated a strong negative correlation (Fig. 1C). In normal human liver sections expression of FoxM1 is low, whereas FoxA2 is expressed at high levels (supplemental Fig. S1B).

**Figure 1:**
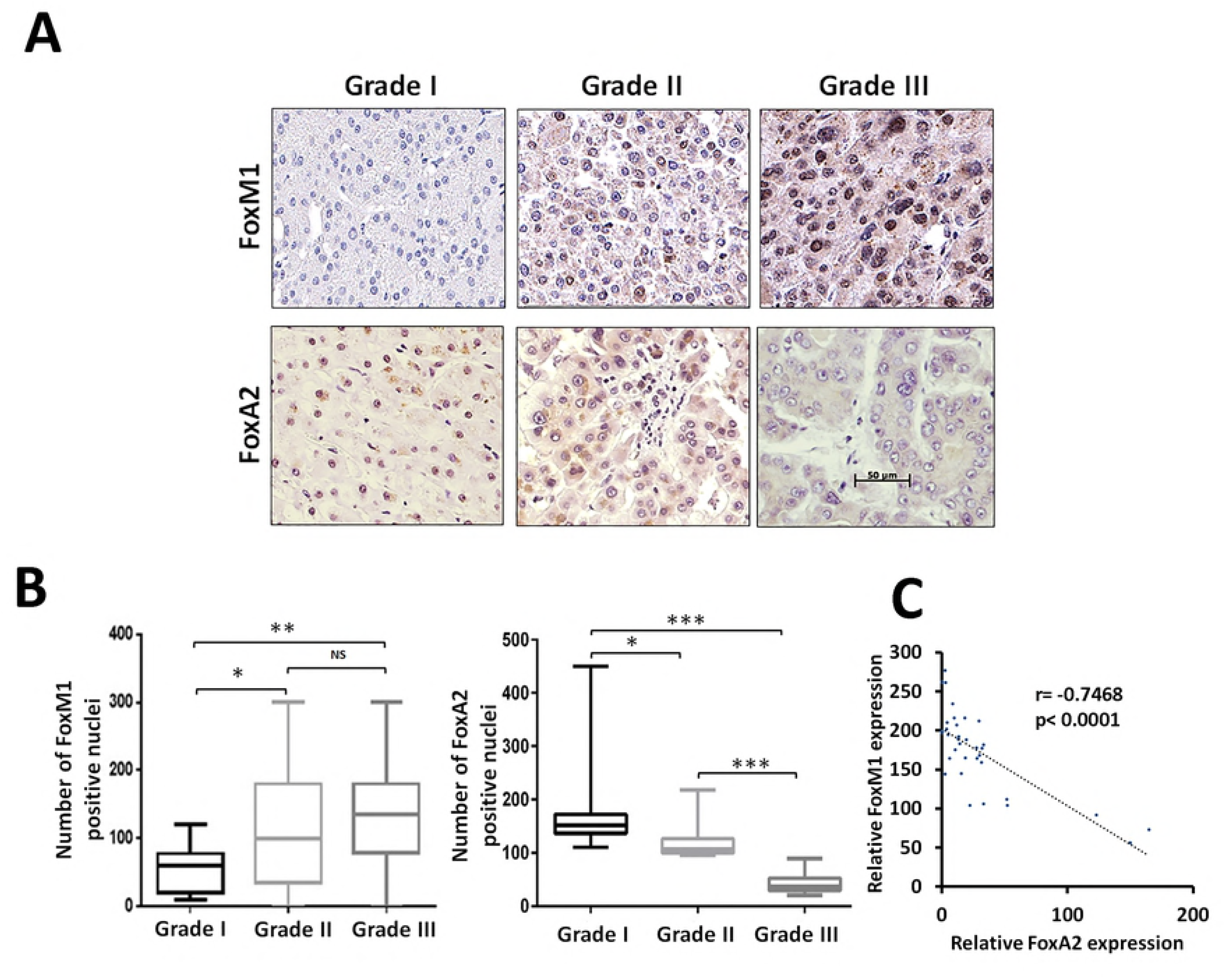
Expression of FoxM1 and FoxA2 in human HCC tumor tissue microarrays: Human HCC tissue microarrays containing grade1 (n= 25), grade 2 (n= 19) and grade 3 (n= 27) were subjected to immunohistochemical staining using FoxM1 and FoxA2 antibodies. (A) The representative tissue cores from grades 1, 2 and 3 are shown. (B) Quantification of the imunohistochemical stains in the tissue cores corresponding to each of the grades was performed. (C) Pearson correlation between FoxM1 and FoxA2 expression is shown. Statistical calculations were performed using GraphPad. All images are in same scale as shown in grade 3 FoxA2 panel.

### Deletion of FoxM1 in a Ras-transgenic model for HCC causes accumulation of FoxA2

Recently, we studied the roles of FoxM1 in HCC progression using a transgenic mouse model that expresses oncogenic HRas in the liver [10]. In that study, the floxed alleles of FoxM1 were deleted after HCC development using the MxCre deletion system, which deletes floxed alleles in liver as well as in blood cells [22], and that somewhat mimics what would be expected from a drug that inhibits FoxM1. In those experiments, FoxM1-deletion in the HCC nodules was detected mainly in the HCC cells [10]. Moreover, we showed that deletion of FoxM1 after HCC development inhibits HCC progression [10]. Sections from those tumor nodules were analyzed for FoxA2 expression by immunohistochemistry. The HRas-derived tumor nodules without FoxM1-deletion exhibited very little expression of FoxA2 (Fig. 2A-B). But, in the FoxM1-deleted samples there was a significant increase in the FoxA2 (Fig. 2A-B). The observation was confirmed by western blot assays using extracts from tumor nodules with and without FoxM1-deletion (Figs. 2C). The increases in the expression of FoxA2 in the FoxM1-deleted samples provides in vivo genetic evidence that FoxM1 plays a role in the inhibition of FoxA2 in HCC. In normal mouse liver FoxM1 is expressed at a low level, whereas the expression of FoxA2 is abundant (supplemental Fig. S1B).

**Figure 2:**
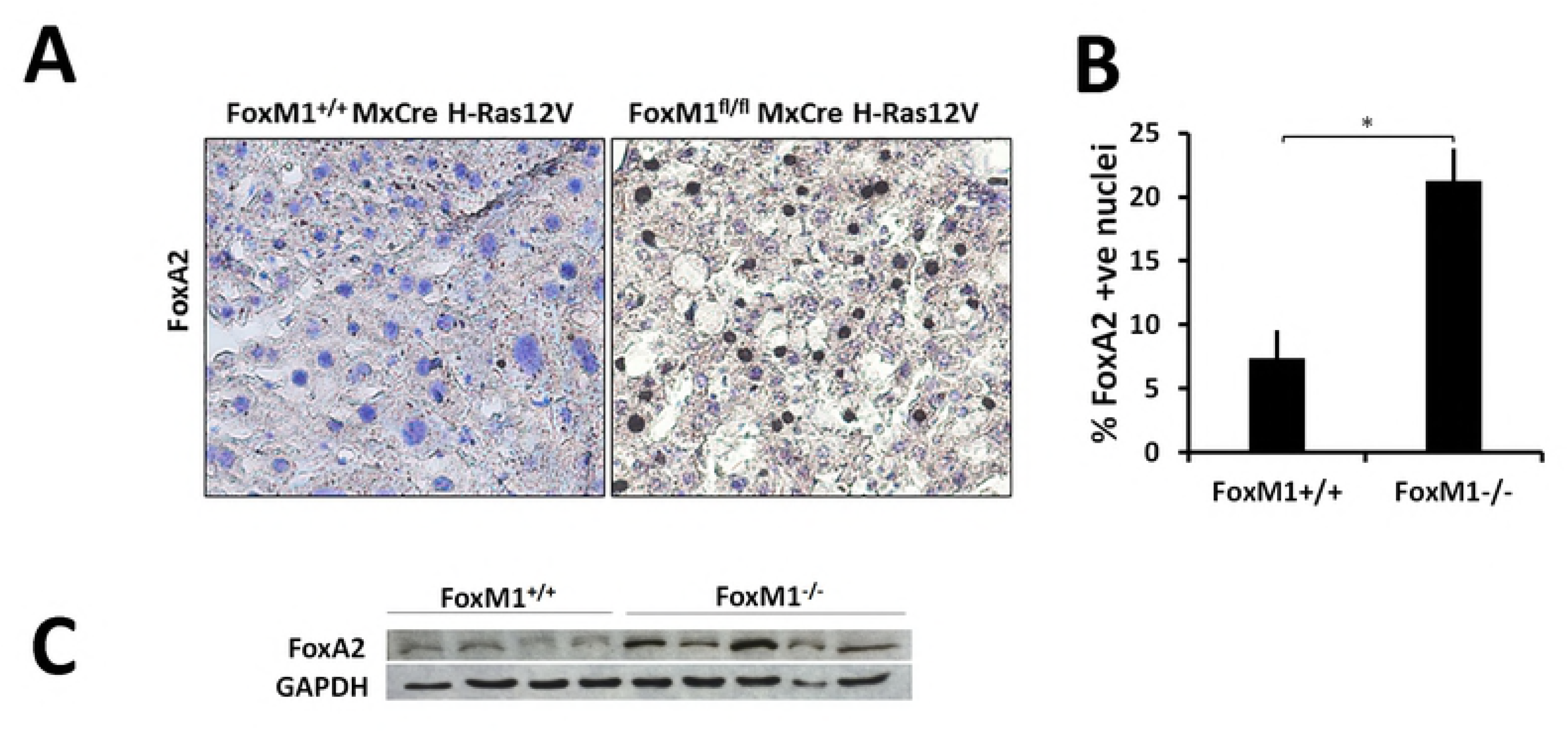
Increase in FoxA2 protein level in mouse HCC samples upon FoxM1 deletion: (A) Immunohistochemical staining of mouse HCC sections using FoxA2 antibody with and without deletion of FoxM1. HCC in those mice was driven by expression of oncogenic H-Ras. The mouse strains also harbored floxed alleles of FoxM1 and MxCre. FoxM1 deletion was induced following HCC development by injecting (5 times) the mice with polyIpolyC. (B) Graphs showing quantification of FoxA2 +ve HCC cells from 3 pairs of mice, and 5 random fields were chosen for analyses. p<0.05 (C) Comparison (0.1 mg of tumor extracts) of the FoxA2 protein-levels between FoxM1+/+ and FoxM1 fl/fl tumor tissue-extracts from mice injected with poly(I)poly(C) five times, using western blots.

### FoxM1 directly inhibits expression of FoxA2 in HCC cells

Next we determined the effects of FoxM1b over-expression and FoxM1-knockdown on the levels of FoxA2 in HCC cell lines. Expression of T7-tagged FoxM1b in Huh7 and HepG2 cells inhibited the levels of FoxA2 at both mRNA (Fig. 3A) and protein levels (Fig. S2A-B). Also, knockdown of FoxM1 in Huh7 and SNU449 cells caused increase in the levels of FoxA2 in both mRNA (Fig. 3B-C) and protein levels (Fig. S2C-D). Also, we developed Huh7 stable cell lines in which FoxM1-shRNA can be expressed in an inducible manner by adding doxycycline in the culture medium. Expression of FoxM1-shRNA in three independent clones increased expression of FoxA2 proteins (Supplemental Fig. S2E).

**Figure 3:**
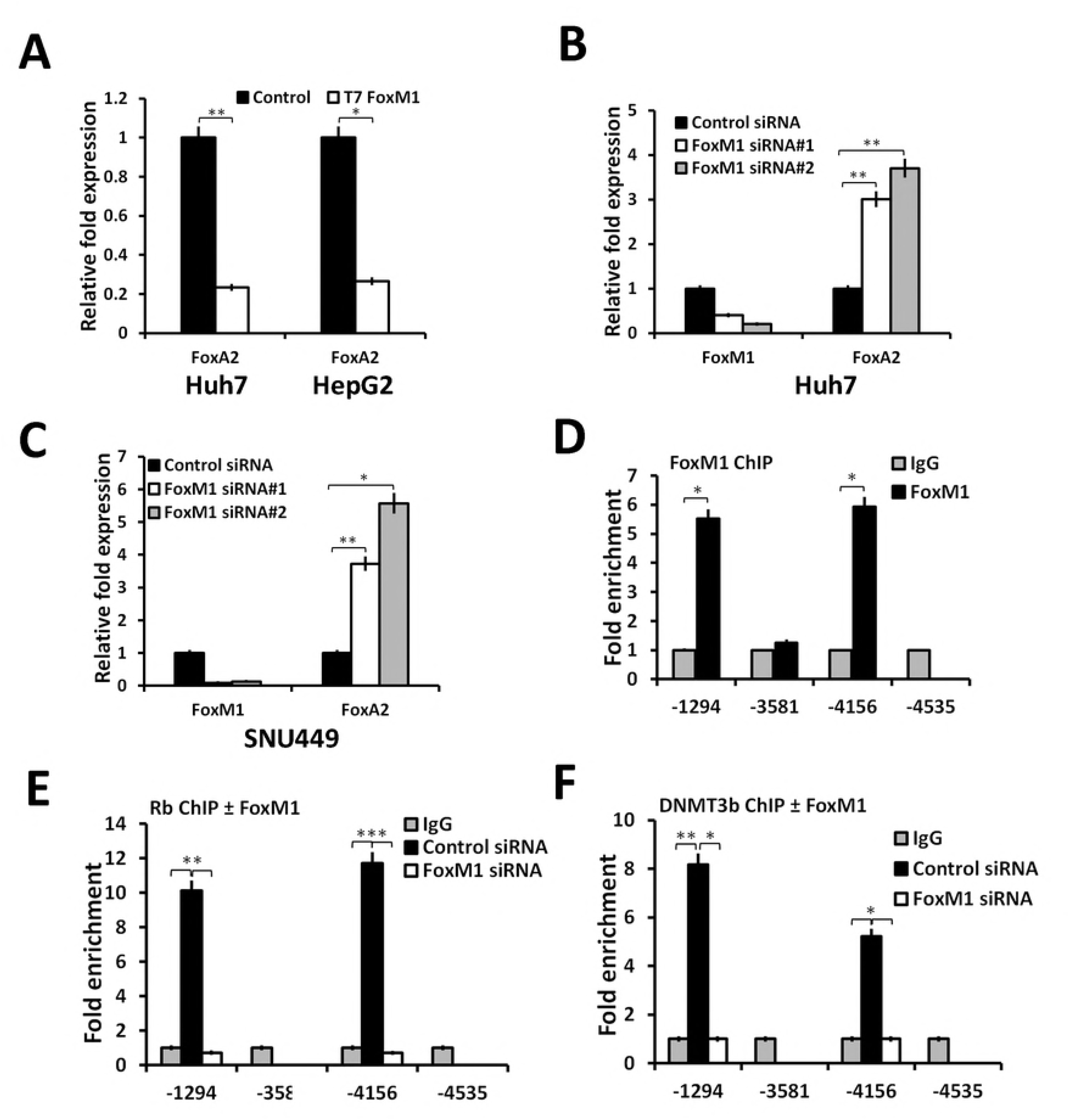
FoxM1 inhibits expression of FoxA2 by recruiting Rb and DNMT3b onto the FoxA2 promoter: (A) Huh7 cells or HepG2 cells were transfected with empty vector (control) or vector expressing T7-FoxM1b. Forty-eight hours after transfection, cells were harvested for RNA assays. Total RNA (1 ug) was analyzed by quantitative real time PCR (qRT-PCR) for expression of FoxA2 and cyclophilin. Huh7 cells (B) or SNU449 cells (C) were transfected with control-siRNA or FoxM1-siRNAs. Seventy-two hours after siRNA transfection, cells were harvested for RNA (1 ug) assays. (D) Huh7 cells were transfected with control-siRNA or FoxM1-siRNA1. Seventy-two hours after transfection, cells were subjected to crosslinking for chromatin-IP (ChIP). The chromatin preparations were immunoprecipitated with FoxM1-ab or IgG. Enrichments of the FoxA2 promoter fragments were assayed by quantitative RT-PCR, and the relative enrichments with FoxM1-ab over that with IgG after normalization against a non-specific site are shown. (E-F) Huh7 cells transfected with control-siRNA or FoxM1 siRNA for 72h, and then processed for ChIP using Rb antibody (E) or DNMT3b-antibody (F). Relative enrichments of the FoxA2 promoter fragments from control-siRNA transfected cells over those from the FoxM1-siRNA transfected cells are shown. Statistical calculations were done using GraphPad Prism online tool for *t*-test and p values stated as *p≤0.05 and **p≤0.001

To determine whether a direct mechanism is in play, we sought to determine whether FoxM1 directly targets the FoxA2 promoter. The human FoxA2 gene contains at least four putative FoxM1-binding elements (Supplemental Fig. S3). Chromatin-IP experiments using FoxM1-ab detected enrichment of DNA fragments encompassing the sites at -1294 and -4156 in the FoxA2 gene (Fig. 3D). The other sites in the FoxA2 upstream regions did not show any significant enrichment over that with the IgG (supplemental Fig. S3B). Previously, we showed that FoxM1 binds to both DNMT3b and Rb, forming a repressor complex in breast cancer cells [19]. Immunoprecipitation of Huh7 cell extracts with a monoclonal antibody against FoxM1 (supplemental Fig. S3C) co-immunoprecipitated DNMT3b and Rb. Therefore, we carried out chromatin-IP experiments with Rb and DNMT3b antibodies using the Huh7 cells expressing control-siRNA or FoxM1-siRNA. Both Rb and DNMT3b bound to the same promoter-fragments that were enriched in chromatin-IP with FoxM1-ab (Fig. 3E-F). Moreover, knockdown of FoxM1 caused significant reduction in the bindings of Rb (Fig. 3E) and DNMT3b (Fig. 3F) onto the specific sites in the FoxA2 promoter. Together, the results confirm the notion that FoxM1 recruits DNMT3b and Rb onto the FoxA2 promoter.

Recruitment of DNMT3b onto the FoxA2 promoter suggests that the repression by FoxM1 would involve methylation of CpG islands. To investigate that, Huh7 cells were transfected with FoxM1b-expresion vector or FoxM1-siRNA. Genomic DNAs from the transfected cells were treated with bisulfite followed by PCR using primers for the CpG islands in the FoxA2 promoter. Expression of FoxM1b caused an increase in CpG methylation near the FoxM1-binding sites in the FoxA2 gene (Fig. 4A). Moreover, knockdown of FoxM1 caused a decrease in CpG methylation at those sites in the FoxA2 promoter (Fig. 4B).

Next, we employed an Rb-shRNA construct that allows inducible depletion of Rb in the presence of doxycycline (Fig. 4C). As shown in Fig. 4D, expression of FoxM1 increased methylation of the FoxA2 promoter in the presence of Rb (no doxycycline), but upon depletion of Rb (doxycycline) there was no increase in the CpG methylation at the indicated sites. Moreover, depletion of Rb caused increases in the expression of FoxA2 (Fig. 4E). These observations demonstrate that the FoxM1/DNMT3b complex methylates and represses the FoxA2 promoters requiring Rb. The extent of FoxA2 repression by FoxM1b varied between 40 and 75%, which is likely due to variations in the levels of active Rb in the transfected cells. Changes in CpG methylation was detected also in the mouse HCC samples following deletion of mouse FoxM1 (Fig. S4A-B) that also binds to Rb and DNMT3b (Fig. S4C).

**Figure 4:**
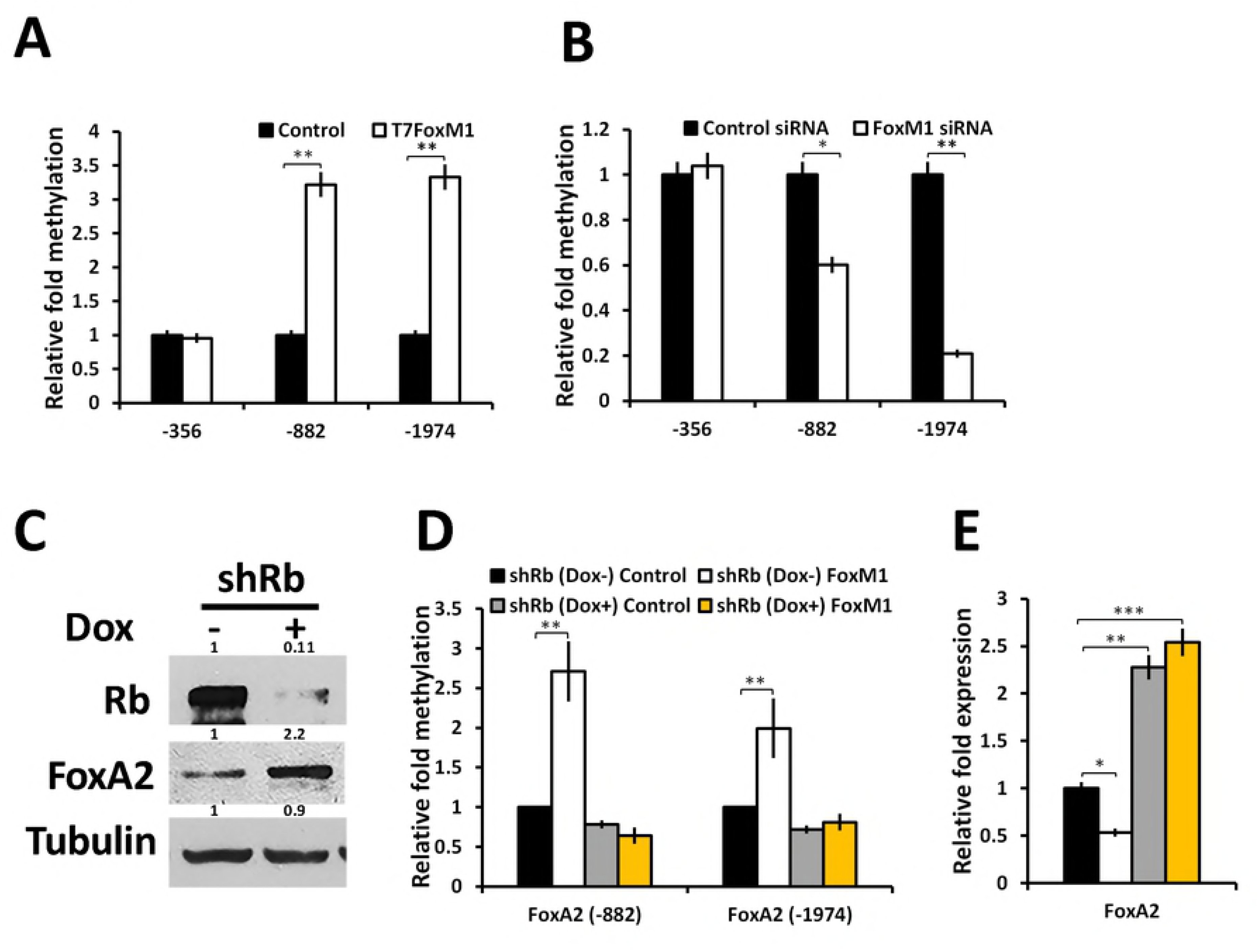
FoxM1 induces methylation of the FoxA2 promoters requiring Rb: (A)Huh7 cells were transfected with empty vector (control) or T7-FoxM1b expressing vector. Genomic DNA was isolated and subjected to bisulphite treatment for CT conversion. Methylation of the FoxA2 upstream regions in control and FoxM1 transfected was assayed using qRT-PCR. (B) Bisulphite converted genomic DNAs isolated from Huh7 cells transfected with control- or FoxM1-siRNA were analyzed for methylation of the FoxA2 promoter by qRT-PCR. (C) Western blot (0.1 mg of cell-extracts) showing depletion of Rb in doxycycline-induced Huh7 cells and effect of Rb knock down on FoxA2 expression. (D) Huh7 cells stably expressing Dox inducible Rb-shRNA were transfected with control or T7-FoxM1 expression vector in the presence or absence of Dox as depicted. The genomic DNAs were isolated for CT conversion using bisulphite method and the difference in the promoter methylation of FoxA2 was assayed by qRT-PCR. (E) Quantification of FoxA2 expression using qRT-PCR in Huh7 cells expressing Dox inducible Rb-shRNA and transfected with either control vector or T7-FoxM1 in absence and presence of Dox. Statistical calculations were done using GraphPad Prism online tool for t-test and p values stated as *p≤0.05and **p≤0.001.

### FoxM1 inhibits FoxA2 in G1 and stimulates pluripotency genes in S/G2/M phases

Given that Rb is required for FoxM1 mediated repression of FoxA2, we predicted that repression occurs mainly in the G1 phase in which Rb is in the underphosphorylated form and is most active. Also, it is the underphosphorylated form of Rb that was shown to associate with FoxM1 [23]. The doxycycline-inducible FoxM1-shRNA cells were treated with vehicle or doxycycline for 96h, and then treated with Hoechst that allows fluorescent staining of DNA in live-cells. The cells were then fractionated using a cell-sorter to obtain cell-populations enriched for G1, S or G2/M cells. As expected expression of the FoxM1-shRNA (Fig. 5A) caused an increase in the G1 population and reductions in the S and G2/M cells (Fig. 5B-D). Clearly, depletion of FoxM1 (open bars in Fig. 5B) caused increases in the expression of FoxA2 in G1 phase. Expressions of the pluripotency genes, on the other hand, were inhibited in the FoxM1-depleted cells, and the inhibition was observed mainly in the S and G2/M cells. c-Myc consistently showed inhibition in the G1 cells. We repeated this experiments with three independent clones of FoxM1-shRNA cells and observed similar results. The changes in gene expression are not indirect effects of G1-inhibition because similar changes were not observed for Oct-4 and Nanog in G1 cells. Also, an unrelated gene Fyn did not show any significant changes in the G1 and S phase cells.

**Figure 5:**
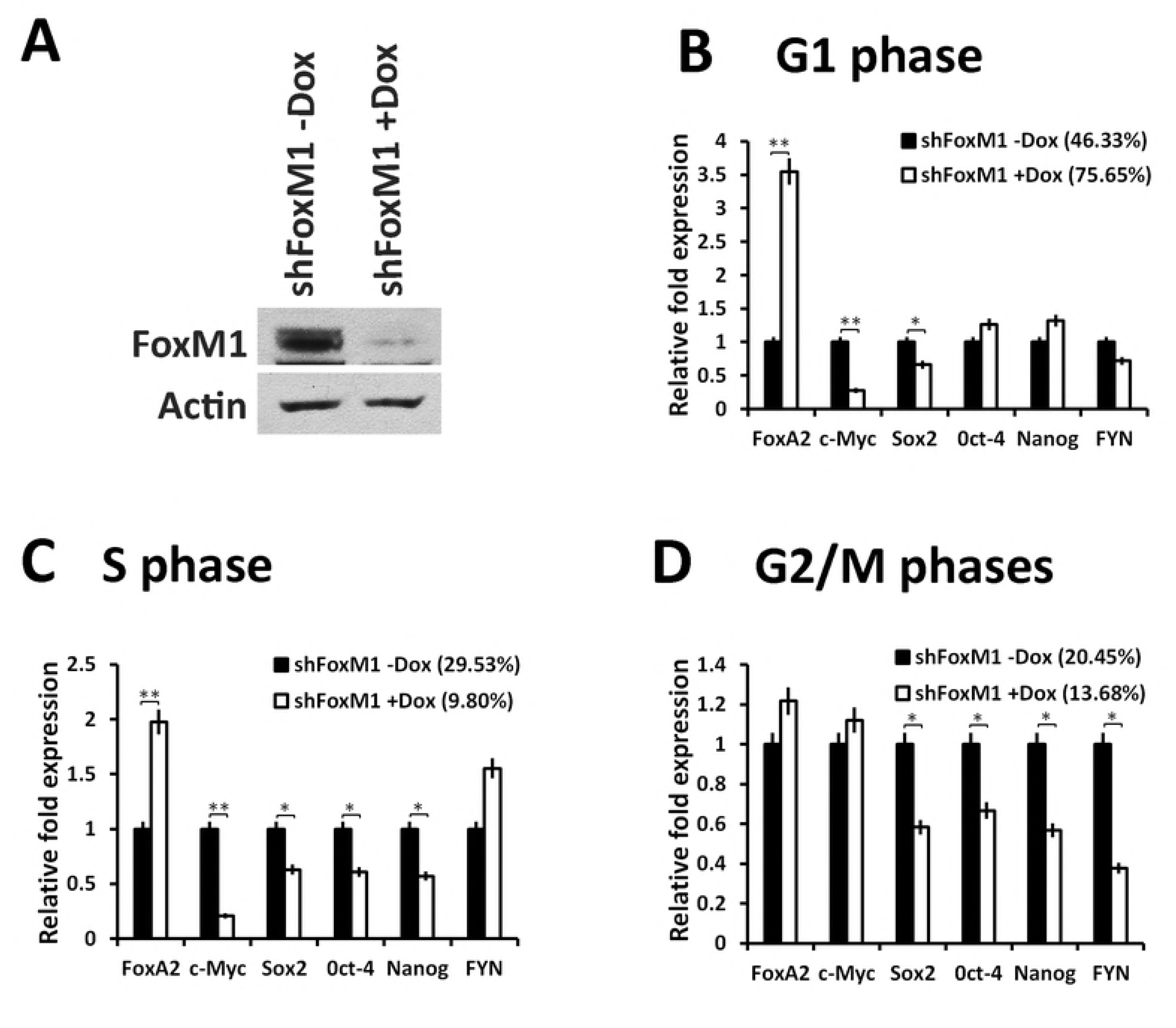
FoxM1 inhibits the expression of FoxA2 in G1 phase of the cell cycle: Doxycycline inducible Huh7-pINDUCER-shFoxM1 cells were either mock treated or treated with 300ng/ml of Doxycycline for 4 days to knock down of FoxM1. Cells were treated with Hoechest 33342 for one hour and then sorted to fractionate cells enriched for G1, S or G2/M phases of the cell cycle. (A) Huh7-pINDUCER-shFoxM1 cells were either mock treated or treated with Doxycycline whole cell extract was prepared at the end of treatment and level of FoxM1 knock down was confirmed by western blotting with FoxM1 antibody and Actin was used as loading control. (B) mRNAs of the indicated genes from G1-phase cells were assayed by quantitative real time PCR. (C) mRNAs of the indicated genes from S-phase cells were assayed by quantitative real time PCR. (D) mRNAs of the indicated genes from G2/M-phases cells were assayed by quantitative real time PCR.

### FoxA2 inhibits the pluripotency genes and blocks auto-activation of FoxM1

Expression of FoxM1 caused a significant increase in the number of spheres when cells were plated in sphere formation media (Fig. 6A-B). Expression of FoxA2, on the other hand, instead of increase the number of sphere, caused decreases in the number of spheres. That is also consistent with the observation that expression of FoxA2 caused inhibition of the pluripotency genes along with FoxM1 and FoxM1 target genes (Fig. 6C and Fig. S5A). Moreover, expression of FoxA2 caused increases in the expression of the hepatocyte differentiation markers ALB, AAT and HNF4α (Fig. 6D). It is noteworthy that inhibition of FoxM1 also increased expression of those differentiation genes (supplemental Fig. S5B).

The inhibition of FoxM1 by FoxA2 is interesting because that could be the mechanism by which FoxA2 inhibits the pluripotency genes. We did not detect an interaction between Rb and FoxA2. Therefore, we considered other possibilities. For example, FoxM1 was shown to auto-activate its own transcription [24]. In chromatin-IP assays we detected interactions of FoxM1 with multiple sites in the FoxM1 promoter (Fig. 6E). Since FoxA2 bind to similar cognate DNA-elements, we considered the possibility that FoxA2 could compete with FoxM1 and inhibit its binding. Consistent with that notion we observed strong inhibitions of FoxM1-binding to its own promoter when FoxA2 was over-expressed (Fig. 6F).

**Figure 6:**
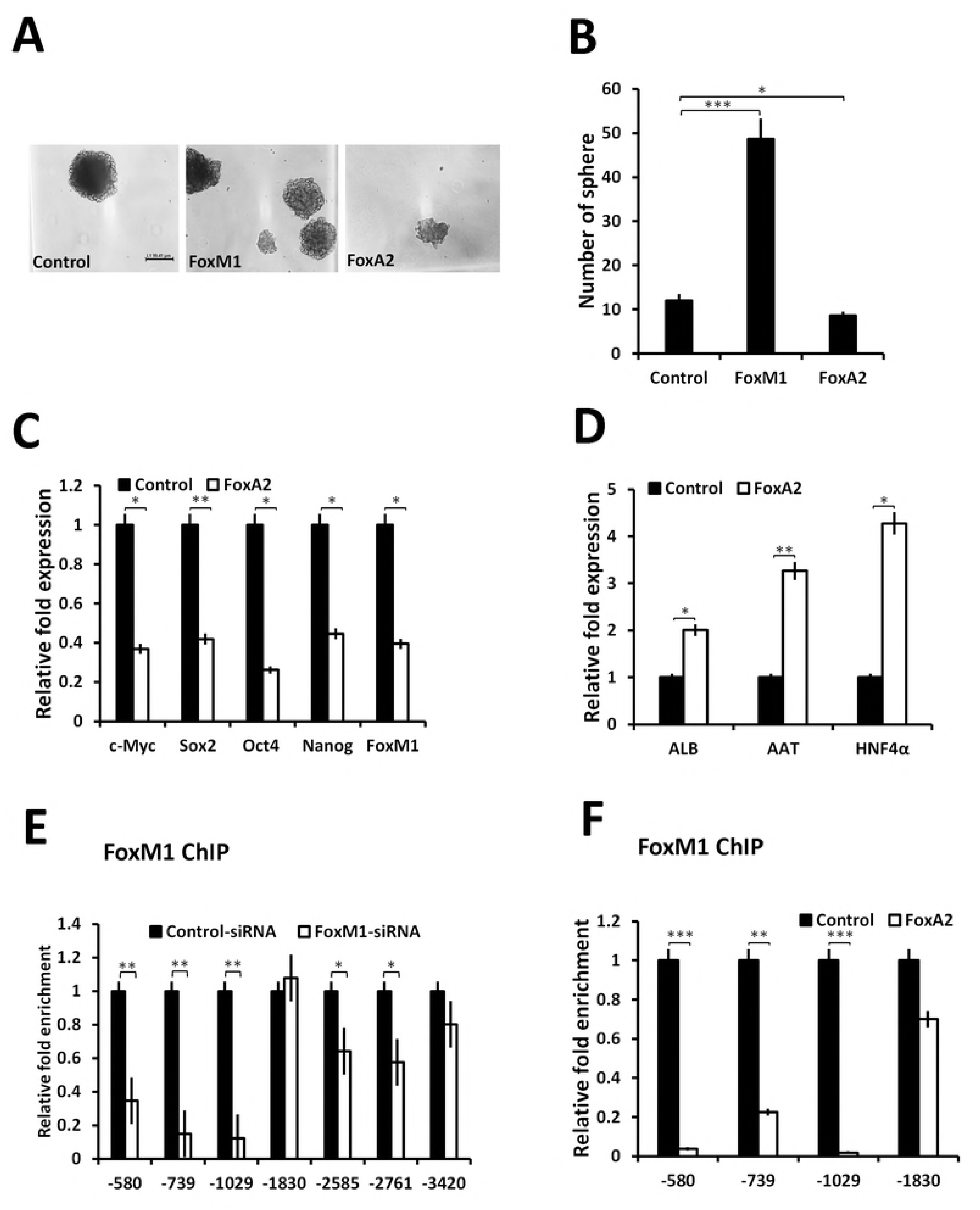
FoxA2 inhibits FoxM1 induced sphere formation by competing for FoxM1 promoter: (A) HepG2 cells stably transfected with empty vector, FoxM1, or FoxA2 expression plasmid as indicated in the figure. Two hundred cells were then seeded in sphere formation medium in low attachment petri dishes. Cells were allowed to grow for a week. Cells were counted and representative picture was captured under microscope. (B) Shows the quantification of sphere formation assay. (C) SNU449 cells were transfected with Empty vector or FoxA2 expression vector. Cells were harvested 48h post-transfection and RNA was isolated to assay for stem cell marker genes and FoxM1 as indicated in figure. (D) Shows the effect of FoxA2 on the indicated liver differentiation genes. (E) SNU449 cell transfected with control siRNA and FoxM1 siRNA were harvested 72h post transfection. Cells were subjected for ChIP assay using FoxM1 antibody. Binding of FoxM1 on FoxM1 promoter were analyzed by qRT-PCR. (F) SNU449 cells were either transfected with control vector or with FoxA2 plasmid. Forty-eight hours post-transfection cells were harvested and processed for ChIP. FoxM1 binding was analyzed using qRT PCR.

### FoxA2 inhibits FoxM1b-induced clonogenicity and soft agar colony formation

FoxM1 is a pro-proliferation transcription factor that also inhibits apoptosis, and drives aggressive progression of cancers when over-expressed [25]. If repression of FoxA2 is important, the prediction is that expression of FoxA2 would inhibit the FoxM1 pathways in HCC cells. We observed that expression of FoxM1 led to significant increases in clonogenicity of the Huh7 cells (Fig. 7A and supplemental Fig. S6A). Expression of FoxA2 alone did not show any significant effect, but when expressed in combination with FoxM1 they strongly inhibited the FoxM1-induced increased clonogenicity of the Huh7 cells (Fig. 7A and supplemental Fig. S6A). Similarly, coexpression of FoxA2 inhibited FoxM1-induced increase in soft agar colonies (Fig. 7B and supplemental Fig. S6B). As expected from the observation that FoxA2 inhibit autoactivation of FoxM1, we consistently observed that expression of FoxA2 affected the levels of total FoxM1. FoxA2 did not have any significant effect on the co-expressed Flag tagged FoxM1b levels (Fig. 7C, Flag panel). However, expression of Flag-FoxM1b increased the levels of the endogenous FoxM1 (Fig. 7C, top panel), and co-expression of FoxA2 inhibited the increase of the endogenous FoxM1 (Fig. 7C, top panel). The results are consistent with a model in which FoxM1 and FoxA2 have opposite regulatory effects on each other.

**Figure 7:**
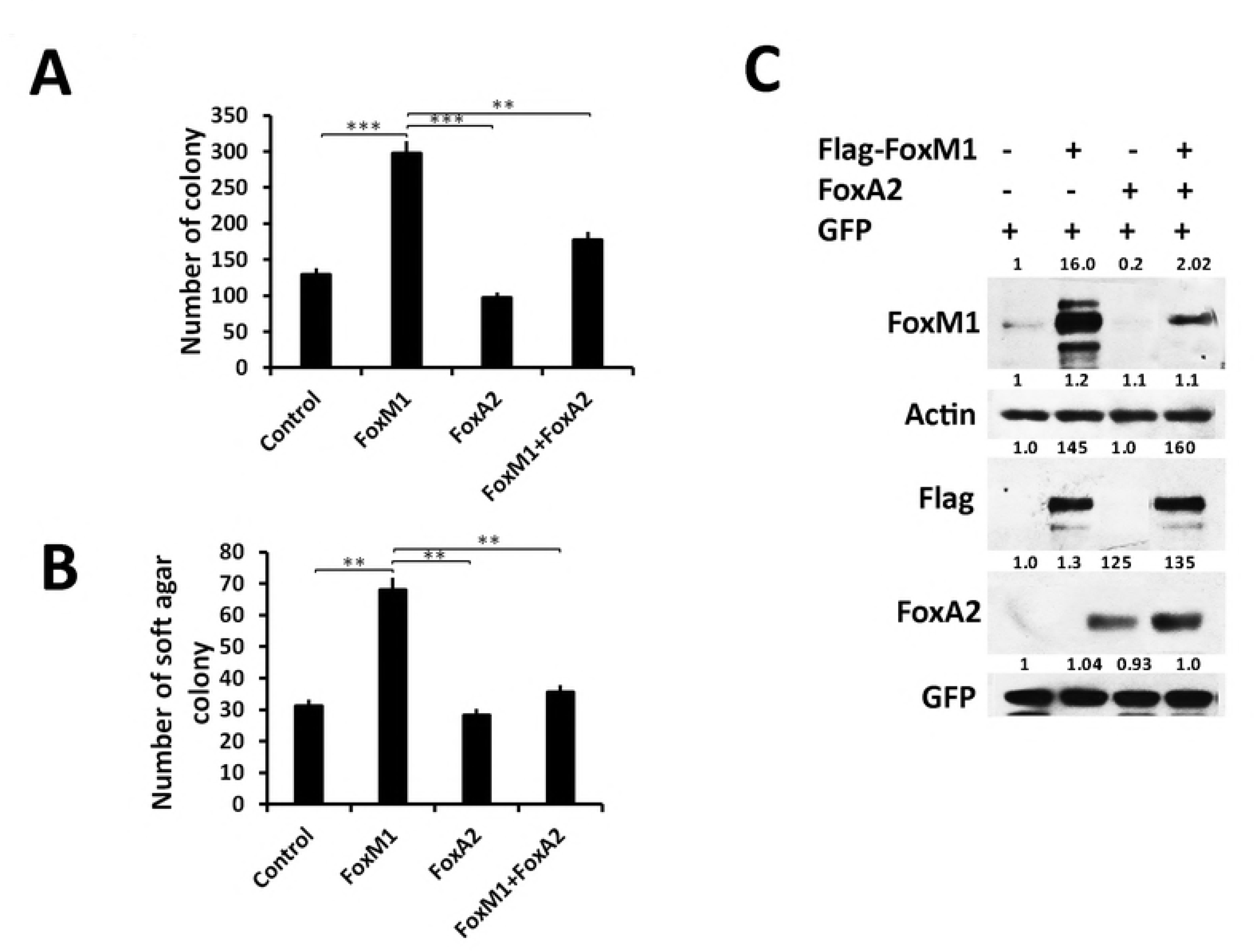
FoxA2 inhibits FoxM1-induced clonogenicity and anchorage independent growth of the Huh7 cells: (A) Huh7 cells were transfected individually with empty vector, Flag-FoxM1 or FoxA2 or a combination of Flag-FoxM1 and FoxA2, along with GFP expression plasmid as transfection control. The transfected cells were subjected to clonogenicity or soft agar assay. About 1×10^4^ cells were seeded in triplicate in 24 well plate and allowed to grow for one week then cells were fixed and stained with crystal violate and pictures (supplemental Fig. S7A) were taken under light microscope. Quantification of the colonies is shown. (B) Huh7 cells (2 x10^4^ cells) as described in panel (A) were seeded into soft agar in triplicate and allowed to grow for 20 days and the representative appearances are shown in supplemental Fig. S7B. Graph shows the quantification of number of distinctly visible colonies with unaided eye. (C) Huh7 cells as described in panel (A) were lysed forty-eight hours after transfection, and 70 ug of extracts were resolved on SDS-PAGE. Expressions of FoxM1 and FoxA2 were assayed by western blotting with anti-Flag, anti-FoxM1, anti-FoxA2 antibodies. For loading control, actin-ab and anti-GFP antibody were used. For the over-expressed Flag-FoxM1b, FoxA2 short exposure (*t*<1 sec) of the blots was taken. Statistical calculations were done using GraphPad Prism online tool for *t*-test and p values stated as *p≤0.05, **p≤0.001 and ***p≤0.0001.

### FoxA2 inhibits FoxM1 and Ras-induced HCC

We showed that FoxM1 is essential for progression of Ras-induced HCC (). Based on the observation that FoxA2 inhibits FoxM1 expression, we predicted that FoxA2 would inhibit Ras-induced HCC progression. We tested that by tail vein injection of FoxA2-expressing adenovirus (Ad-FoxA2) in 10-months old Alb-Hras12V transgenic mice harboring HCC. Five male mice were injected with Ad-FoxA2 or Ad-LacZ twice. The second injection was done 12-days after the first and the livers were harvested 10-days after the second injection. Expression of FoxA2 caused significant reductions in the number of tumor nodules and tumor burden. Representative livers with HCC nodules are shown in Fig. 8A, and quantifications are shown in Fig. 8B-C. The inhibition of HCC progression was associated with inhibition of FoxM1 expression as well as Ki67+ve cells (Fig. 8D-F). These observations provide in vivo evidence that FoxA2 inhibits FoxM1 and Ras-induced HCC progression.

**Fig 8:**
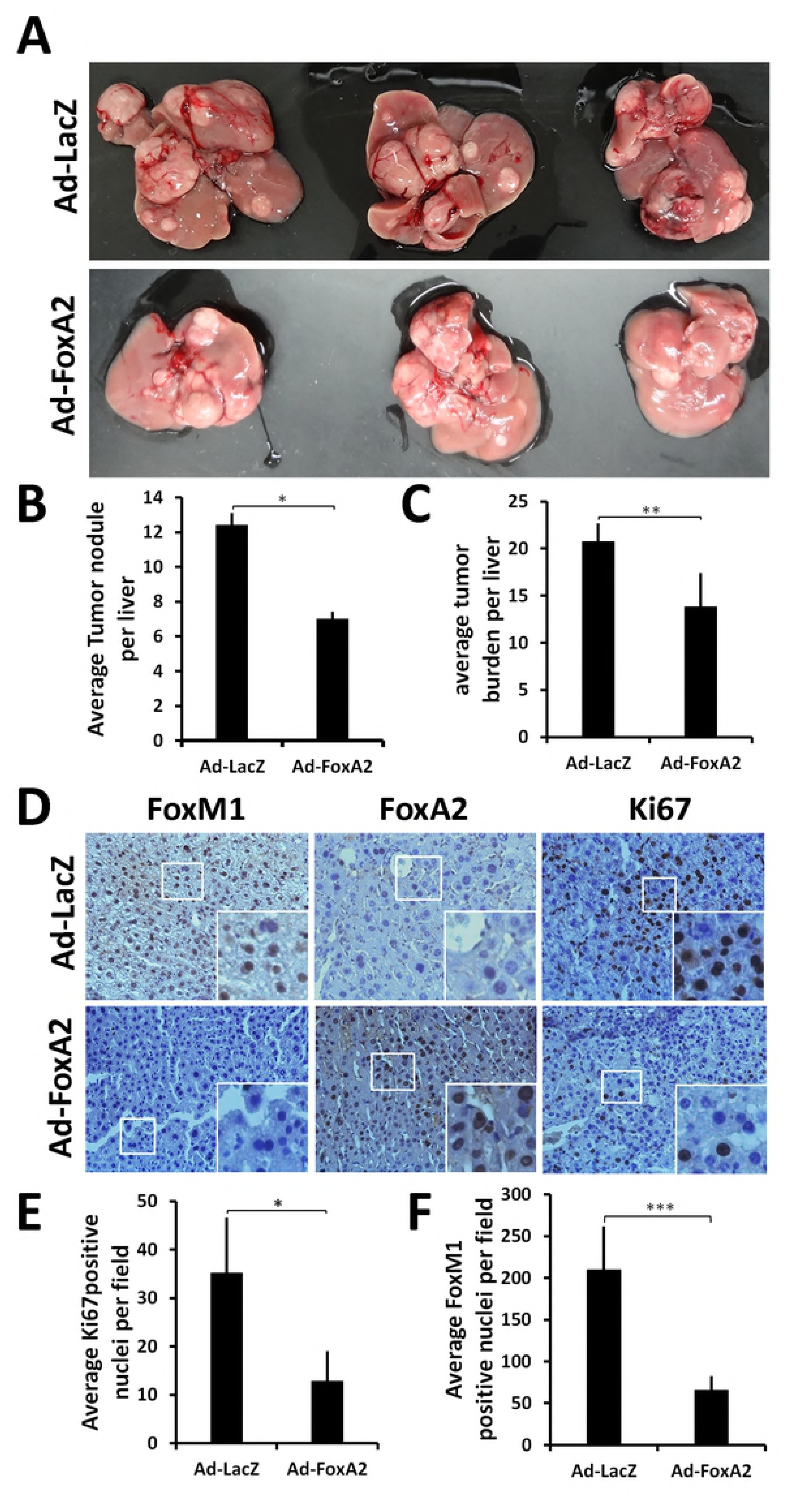
Adenoviral Foxa2 expression inhibits HCC progression and FoxM1 expression. Alb-H-Ras12V male mice (5 per group) at 10 months of age were injected via the tail vein with purified Ad-FoxA2 or Ad-LacZ (0.2 ml at 1.7X10^10 pfu per ml). Twelve days following first injection the mice were injected again with the same virus and sacrificed after 10 days. (A) Livers from three mice of each group are shown. Average number of tumor nodules (B) and tumor burden (C) are plotted. (D) IHC staining for FoxM1, FoxA2 and Ki67 are shown. Average Ki67+ve nuclei/field (E) and average FoxM1+ve nuclei/field from 20 different fields of sections from 5 mice are plotted. Statistical calculations were done using GraphPad Prism online tool for *t*-test and p values stated as *p≤0.05, **p≤0.001 and ***p≤0.0001.

## DISCUSSION

Results presented here are significant in several ways. First, our observations provide insights into the mechanisms by which FoxM1 drives accumulation of poorly differentiated cancer cells during high-grade progression of HCC. We show that FoxM1 suppresses expression of the hepatocyte-differentiation genes FoxA2 in G1 phase, and that suppression is important for expression of the pluripotency genes in S/G2 phases of the cell cycle. Also, we show that FoxA2 regulates FoxM1 expression by blocking its auto-activation mechanism. Our results suggest that it is the level of FoxM1 versus FoxA2 that determine the differentiation state of the HCC cells.

Reduced expression of FoxA2 coincides with over-expression of FoxM1 (Fig. 1). The need for over-expression of FoxM1 might be related to low-abundance of active Rb in HCC cells. High levels of FoxM1 would be needed to seek out active under-phosphorylated Rb protein because it is the under-phosphorylated Rb that binds to FoxM1 [23]. We suspect that the FoxM1 mediated inhibition of FoxA2 might be involved in de-differentiation of the HCC cells or maintenance of poorly differentiated HCC cells. Alternatively, in the event that the low- and high-grade HCCs develop from different progenitors, we speculate that the FoxM1 mediated repression of FoxA2 is important for the progenitors that give rise to high-grade HCC.

The involvement of Rb in the suppression of FoxA2 also suggests that the mechanism might be more active in G1 phase where under-phosphorylated Rb is abundant. Consistent with that, depletion of FoxM1 caused accumulation of FoxA2 mainly in G1 phase. Interestingly, studies on embryonic stem (ES) cell differentiation indicated that G1 phase is the phase in which the chromatin is available for the differentiation mechanisms, and that the ES cells retain pluripotency by suppressing differentiation mechanisms in G1 and increasing expression of the pluripotency genes in the S/G2 phases [26]. In the light of those observations in ES cells, our observations that FoxA2 inhibits expression of the pluripotency genes and that FoxM1/Rb/DNMT3b inhibit FoxA2 are interesting because they explain how FoxM1 over-expression maintain poorly differentiated state of cells in high-grade HCC. The involvement of Rb in the suppression of the FoxA2 gene is surprising because it suggests that Rb participates in progression of high-grade HCC. We show that FoxM1 recruits Rb and DNMT3b onto the promoter of FoxA2, and increases methylation of the CpG islands in those promoters. Moreover, depletion of Rb blocks FoxM1-mediated increase in promoter-methylation and suppression of the FoxA2 gene. Together, the results suggest that, in the context of over-expressed FoxM1, there is a gain of function for Rb, and that function is related to the suppression of differentiation genes, which is likely involved in high-grade progression of HCC.

Inhibition of FoxM1-induced increase in clonogenicity and soft-agar colony formation by FoxA2 further explains why expression of FoxA2 needs to be repressed by FoxM1. Moreover, FoxA2 inhibits FoxM1 in Ras-induced HCC. These observations provide evidence for a new regulatory loop in which over-expression of FoxA2 inhibits expression of FoxM1 and the FoxM1 target genes in HCC cells. FoxM1 was shown to activate its own expression in a positive feedback loop [24]. We think that FoxA2 disrupts that positive feedback, resulting in an inhibition of FoxM1 expression. That would further explain the opposite expression patterns of FoxM1 and FoxA2 in HCC.

## EXPERIMENTAL PROCEDURES

### Cell Culture and Transfections

Human hepatocellular carcinoma Huh7 cells (American Type Culture Collection) were maintained in Dulbecco’s modified Eagle’s medium supplemented with 10% fetal bovine serum (HyClone Laboratories Inc.) for normal cell culture and Tet System Approved FBS (Clontech, Mountain View, CA, Cat No. 631105) for inducible cell lines with 100 units of penicillin/ streptomycin at 37°C with 5% CO2. Cells were transfected with plasmid DNA or siRNA using Lipofectamine^™^ 2000 (Invitrogen) in serum-free tissue culture medium following the manufacturer’s protocol. Six hours after transfection, cells were fed with complete Dulbecco’s modified Eagle’s medium containing 10% fetal bovine serum.

### Immunohistochemistry and tissue microarray

Immunohistochemical stainings were performed following standard procedure. Antigen retrieval was done using sodium citrate buffer and sections were then treated with antibodies overnight. Additional blocking step was performed using Avidin/biotin Vectastain kit following manufacturer’s protocol. Visualization was done using DAB and counterstained using Hematoxylin (Polyscientific, Bay Shore, NY). For antibodies of mouse origin, mouse on mouse (MOM) kit was used. All used reagents are from Vector Labs (Burlingame, CA). Information about the antibodies is included in supplemental table 1.

### Animal studies

All animal experiments were pre-approved by the UIC institutional animal care and use committee. Previously described Alb-H-ras12V mice were crossed with FoxM1fl/fl MxCre C57/BL6 mice to obtain FoxM1fl/fl MxCre Alb-H-ras12V and FoxM1+/+ MxCre Alb-H-ras12V mice. For deletion studies, eight months old male mice (FoxM1+/+ MxCre Alb-H-ras12V and FoxM1fl/fl MxCre Alb-H-ras12V) were subjected to five or ten intraperitoneal (IP) injections (every other day) with 250μg of synthetic polyinosinic-polycytidylic acid (polyIpolyC) (Sigma-Aldrich, St. Louis, MO) to induce expression of the Mx-Cre transgene. The mice were sacrificed three weeks following the last injection, and the liver tissues and HCC nodules were harvested. For FoxA2 expression in mouse liver, tail vein injections (30 gauge needle) with 0.2 ml of adenovirus expressing FoxA2 or LacZ (1.7X10^10^ pfu/ml) in 10 months old Alb-H-Ras12V male mice were carried out under anesthesia.

### RT-PCR, Western Blot, and Chromatin Immunoprecipitation

RNA was Trizol extracted (Invitrogen, Carlsbad, CA) and cDNA was synthesized using Bio-Rad reverse transcriptase (Bio-Rad, Hercules, CA). cDNA was amplified using SYBR Green (Bio-Rad, Hercules, CA) and analyzed via iCycler software. Western blots and chromatin-IPs were performed following previously described procedures [19]. For chromatin-IPs, signals obtained with IgG and specific-antibodies were first normalized with signals obtained with those antibodies on a non-specific site in the GATA3 promoter [19]. The normalized values were used to plot the fold enrichment with FoxM1-ab over IgG. For Rb-ChIP and DNMT3b-ChIP, we compared enrichments with same antibody, after normalization, in the presence and absence of FoxM1-siRNA, and the fold enrichments in control-siRNA over FoxM1-siRNA were plotted. All primer sequences and antibodies are included in supplemental table 1.

### Isolation of genomic DNA

Genomic DNA (gDNA) was obtained from Huh7 cells and mouse tissue using Wizard™ Genomic DNA Purification kit as instructed by the manufacturers manual.

### Bisulfite treatment and Quantitative methylation-specific PCR assay (qMSP)

Genomic DNA samples were treated with EZ DNA methylation™ kit (Zymo Research, Orange, CA) according to the manufacturer’s recommendation. The extent of methylation of desired gene was then measured by qPCR amplification with pairs of specific primers as mentioned in supplemental table-1 which were designed using MethPrimer MSP/BSP prediction and primer designing tool. Quantitative MSP was performed with using SYBR Green (Bio-Rad, Hercules, CA) and analyzed via iCycler software. Each reaction contained 20 ng of bisulfite-treated DNA as a template, 6.25 μl SYBR Green PCR (Bio-Rad, Hercules, CA) and 50 nM each forward and reverse primers in a total volume of 12.5 μl. The quantification cycle (C_q_) was determined for each reaction with methylation-specific primers (MSP) the ratio of unmethylated to total amplifiable bisulfite-treated DNA was calculated.

### Statistical analysis

Statistical significance was calculated by the Student’s *t* test [27] and Pearson correlation coefficient. Statistically significant changes were indicated with asterisks (*** p <0.001; **p<0.01;*p<0.05).

## AUTHOR CONTRIBUTIONS

VC and AP carried out the experiments. GG generated and provided the human HCC tumor microarrays. DK helped designing the mouse experiments. PR was involved in designing the experiments and writing the manuscript.

## ACKNOWLEDGMENT

Authors thank Dr. S. Elledge (Harvard Medical School, Boston) for sharing the Rb-shRNA construct. The work was supported by grants from National Institute of Health (CA 177655 and CA 175380) to PR. PR is supported also by a Merit Review Grant (BX000131) from the Veteran Administration.

